# Structural, *in silico*, and functional analysis of a Disabled-2-derived peptide for recognition of sulfatides

**DOI:** 10.1101/2020.05.06.081299

**Authors:** Wei Song, Carter J. Gottschalk, Tuo-Xian Tang, Andrew Biscardi, Jeffrey F. Ellena, Carla V. Finkielstein, Anne M. Brown, Daniel G. S. Capelluto

## Abstract

Disabled-2 (Dab2) is an adaptor protein that regulates numerous cellular processes. Among them, Dab2 modulates the extent of platelet aggregation by two mechanisms. In the first mechanism, Dab2 intracellularly downregulates the integrin α_IIb_β_3_ receptor, converting it to a low affinity state for adhesion and aggregation processes. In the second mechanism, Dab2 is released extracellularly and interacts with both the integrin α_IIb_β_3_ receptor and sulfatides, both of which are known to be pro-aggregatory mediators, blocking their association to fibrinogen and P-selectin, respectively. Our previous research indicated that a 35-amino acid region within Dab2, which we refer to as the sulfatide-binding peptide (SBP), contains two potential sulfatide-binding motifs represented by two consecutive polybasic regions. Using a combined methodology including molecular docking, nuclear magnetic resonance, lipid-binding assays, and surface plasmon resonance, this work identifies the critical Dab2 residues within SBP that are responsible for sulfatide binding. A hydrophilic region, primarily mediated by R42, is responsible for the interaction with the sulfatide headgroup, whereas the C-terminal polybasic region contributes to interactions with the acyl chains. Furthermore, we demonstrated that, in Dab2 SBP, R42 significantly contributes to the inhibition of platelet P-selectin surface expression. The interacting Dab2 SBP residues with sulfatide resemble those described for sphingolipid-binding in other proteins, suggesting that sulfatide-binding proteins share common binding mechanisms.

## INTRODUCTION

The adaptor protein Disabled-2 (Dab2) is a multimodular signaling molecule involved in a variety of cellular processes including protein trafficking, cell growth and differentiation, cell adhesion, and modulation of platelet aggregation (1). Two Dab orthologs, Dab1 and Dab2, are present in mammals. Dab1 is primarily expressed in the brain (2), whereas Dab2 is ubiquitously expressed in different tissues (3, 4). Dab2 expression levels have been reported to exhibit a significant effect on cancer initiation and progression. Indeed, Dab2 expression is lost in breast, ovarian, prostate (5), bladder (6), and colorectal cancer cells (7), suggesting that Dab2 can act as a tumor suppressor (8). Two alternative spliced forms of Dab2 are expressed in humans, p96 and p67, with the latter lacking a central exon (3). Dab2 contains a phosphotyrosine-binding (PTB) domain located at the N-terminus, clathrin-binding sites, NPF and DPF motifs, and a proline-rich SH3 domain located at the C-terminus, indicating that Dab2 primarily functions as an adaptor protein.

Recently, Dab2 was characterized as a negative regulator of platelet aggregation by modulating both pro-aggregatory inside-out and outside-in signaling pathways (1). Modulation of the inside-out signaling by Dab2 is mediated by its cytosolic S24-phosphorylated form, which binds to the β3 subunit of the integrin α_IIb_β_3_ receptor, downregulating fibrinogen-mediated adhesion (9). In outside-in signaling, ligand-receptor complexes on activated platelets promote platelet spreading, granule secretion, stabilization of the adhesion and aggregation of platelets, and clot retraction (10). Some of the components of the intracellular granules are required to limit the extent of platelet activation, adhesion, and aggregation. Platelet function must be tightly modulated, as uncontrolled platelet activation can trigger unwanted clinical complications such as thrombocytopenia and thrombosis (11). Dab2 is released from α-granules and relocates to the platelet surface where it modulates outside-in signaling (12). To do this, Dab2 presents two extracellular targets. At the platelet membrane surface, Dab2 associates with the extracellular domain of the α_IIb_ subunit of the integrin receptor *via* its RGD motif, preventing integrin-fibrinogen interactions. Dab2 also binds to sulfatides (13), a class of negatively charged sphingolipids found on most eukaryotic cell surfaces (14). Interestingly, cleavage of Dab2 by the platelet agonist thrombin may be prevented when Dab2 is associated with the sphingolipid (13). Binding of Dab2 to sulfatides limits P-selectin association to sulfatides (15), which is required to prolong platelet aggregation (16).

Initial studies reported that Dab2 binds sulfatides at its N-terminus including the PTB domain (N-PTB; residues 1-241 in human Dab2) (13). Combined mutations in residues K25, K49, K51, and K53 markedly reduce sulfatide binding and platelet aggregation (13). Further studies showed that an α-helical-rich region of 35 amino acids within the Dab2 N-PTB, which was predicted to contain two potential sulfatide-binding motifs (referred to as the sulfatide-binding peptide, SBP), mimic the inhibitory platelet-platelet interaction effects of Dab2 N-PTB (17). In this report, by using a combination of structural, molecular docking, and functional approaches, we refined the mode by which SBP, the minimal sulfatide-binding unit derived from Dab2, associates with sulfatides. We found that the Dab2 SBP residues around α-helix 1 are required for interactions with the sulfatide head group, whereas helix 2 mediates sulfatide acyl chain interactions. Furthermore, we predict that R42, located in the first α-helix of Dab2 SBP, forms hydrogen bonds with the OS atoms of the sulfate group in the sulfatide. We show that both the charge and stereochemistry of R42 is critical for Dab2 SBP’s sulfatide recognition and inhibition of platelet P-selectin surface expression. Results from this report provide details of how Dab2 interacts with sulfatides, which can be used for the design of a Dab2-derived peptide that can block sulfatide interactions, and, consequently, prevent undesired platelet aggregation events.

## MATERIAL AND METHODS

### Chemicals

The list of chemicals and their suppliers are: brain sulfatides, 1,2-dipalmitoyl-sn-glycero-3-phosphocholine (DPPC), 1,2-dipalmitoyl-sn-glycero-3-phosphoethanolamine (DPPE) (Avanti-Lipids), N-dodecylphosphocholine (DPC) (Anatrace), cholesterol (Sigma), and isopropyl β-D-thiogalactopyranoside (IPTG) (Research Products International). All other chemicals were of analytical grade.

### Expression and purification of Dab2 SBP

A cDNA, representing residues 24-58 in human Dab2, was cloned into a pGEX6P1 vector (GE Healthcare). A glutathione S-transferase (GST) fusion and untagged Dab2 SBP were expressed and purified as described (18). Site-directed mutagenesis of Dab2 SBP was carried out using the QuikChange (Stratagene) protocol.

### Creation and validation of Dab2 SBP models

Energy minimized Dab2 SBP and its mutant structures were created with Schrödinger-Maestro v2018-3 (19), using the NMR-derived, Dab2 SBP DPC-micelle embedded structure [PDB ID: 2LSW (17)] as a template. To align with the experimental peptide sequence in this work, residues 19-23 (GPLGS) were built onto the N-terminus of PDB ID: 2LSW using Schrödinger-Maestro’s 3D-Builder. Following the addition of residues 19-23 to the template structure, energy minimization was performed using the OPLS3e forcefield (20) and default settings in Schrödinger-Maestro v2018-3. After energy minimization, the Dab2 SBP structure was exported into PDB format and model quality was assessed using several validation methods. Ramachandran plots generated using the Rampage webserver (21) assessed model quality with respect to favorable and allowed φ and ψ angles of residues, and SWISS-MODEL (22, 23) calculated a QMEAN plot to quantify model quality with respect to solvation and torsion angle. ProSA (24, 25) compared the model to experimentally derived structures similar in composition. Five mutant peptides were individually created using the energy minimized Dab2 SBP structure as the starting template. Each mutation (Y38A, R42A, R42K, K49A/K51A/K53A, and Y50A) was built using Schrödinger-Maestro (19) by altering the residue(s) of interest using the built-in mutagenesis tool. Following residue alteration, energy minimization was performed as stated above. Structures were exported as PDB files and model quality was assessed using the same procedure as Dab2 SBP, with varying validation quality as influenced by the point mutations.

### Molecular docking of Dab2 SBP variants to sulfatide

The structure of sulfatide was obtained from the ZINC database (ZINC96006133) and converted from the mol2 to the PDB format using UCSF Chimera (26). Sulfatide and Dab2 SBP and point mutation structures were prepared in AutoDockTools v1.5.6 (27) and sulfatide was docked to each Dab2 SBP energy minimized structure using AutoDock Vina v1.1.2 (28). The same grid box and center was used for each docking experiment and covered the entire Dab2 SBP structure. The center coordinates for the grid box (X, Y, Z) were (−1.722, 0.667, −2.2194). The grid box dimensions (X, Y, Z) were (30Å, 46Å, 42Å), using a 1 Å grid space. Nine poses were created from each docking run, with the lowest energy pose used for further analysis on residue interaction and fingerprint analysis. Molecular visualization of docked poses was performed in PyMOL (29). Sulfatide pose volume occupancy was visualized using UCSF Chimera (26). Schrödinger-Maestro’s Interaction Fingerprints analysis tool was used with default parameters on each lowest energy docked pose to determine potential peptide-sulfatide interactions. The output matrix was converted manually to a table format and organized by interaction type.

### NMR spectroscopy

^15^N-labeled Dab2 SBP (1 mM) was prepared in 90% H_2_O, 10% ^2^H_2_O, 10 mM *d*_*4*_-citrate (pH 5.0), 40 mM KCl, 1 mM NaN_3_, and 200 mM *d*_*38*_-DPC in the absence or presence of 8 mM sulfatide. A Bruker Avance III 800 spectrometer with cryoprobe and standard Bruker two dimensional ^1^H–^15^N HSQC-type pulse sequences were used to obtain ^15^N longitudinal (*R*_1_) and transverse (*R*_2_) relaxation rates, and ^1^H–^15^N NOEs of Dab2 SBP in the absence and presence of sulfatide at 25°C. Relaxation delays for the *R*_*1*_ experiments were 5, 50, 150, 250, 400, 550, 750, 1200, and 2000 ms. Relaxation delays for the *R*_*2*_ experiments were 17, 34, 51, 68, 119, 153, 203, and 254 ms. The recycle delay for *R*_*1*_ and *R*_*2*_ experiments was 2 s. ^1^H–^15^N NOEs were measured by comparing intensities of ^1^H–^15^N correlation spectra with either 5 s of ^1^H saturation or a 5 s delay preceding ^1^H–^15^N correlation.

### Lipid-protein overlay assay

Sulfatide strips were prepared by spotting 1 μl of the indicated amount of sulfatides, dissolved in chloroform/methanol/water (1:2:0.8), onto a Hybond-C extra membranes (GE Healthcare). Membrane strips were blocked with 3% fatty acid-free BSA (Sigma) in 20 mM Tris-HCl (pH 8.0), 150 mM NaCl, and 0.1% Tween-20 for 1 h at room temperature. Then, membrane strips were incubated with the indicated GST fusion peptide in the same buffer overnight at 4°C. Following four washes with the same buffer, bound fusion proteins were detected with rabbit anti-GST antibody (Proteintech) and donkey anti-rabbit-horseradish peroxidase antibody (GE Healthcare). Binding of fusion peptides to sulfatides was probed using the Supersignal West Pico chemiluminescent reagent (Pierce).

### Surface plasmon resonance

Binding of Dab2 SBP peptides to sulfatide liposomes were monitored by surface plasmon resonance (SPR) at room temperature using a BIAcore X-100 instrument (GE Healthcare). One-hundred nanometer size-calibrated liposomes were generated as previously described (13). Briefly, lipids including DPPC:DPPE:cholesterol (1:1:1; control), or DPPC:DPPE:cholesterol:sulfatide (1:1:1:4), were solubilized in chloroform/methanol/water mixtures. The resulting mixture was dried under N_2_ followed by vacuum to remove solvent traces. Lipid mixture was then resuspended at 0.8 mg/ml in 20 mM Tris-HCl (pH 6.8) and 100 mM NaCl to reach a final concentration of 0.5 mM sulfatide, sonicated, and extruded for 100-nm liposome size at 68°C. L1 sensorchips were first equilibrated with 20 mM Tris-HCl (pH 7.4) and 100 mM NaCl as a running buffer. Then, L1 sensorchips were pretreated with 40 mM octyl β-D-glucopyranoside. At a 30 ml/min flow rate, control liposomes were immobilized onto one of the L1 sensorchip channels whereas a second channel was loaded with sulfatide-containing liposomes. Typical liposome loading was ~4,000 response units/sensorchip channel. The apparent *K*_D_ values were estimated using the BIAevaluation software, version 2.0 (GE Healthcare).

### Blood collection and platelet purification

Blood samples were obtained from healthy volunteers by venipuncture, according to Virginia Tech Institutional Review Board guidelines. Blood samples were collected into vacutainer ACD solution A blood tubes (Becton, Dickinson and Co) and centrifuged for 20 min at 200xg to separate a platelet-rich plasma (PRP) fraction from erythrocytes. Then, the PRP fraction was further centrifuged for 5 min at 1,100x*g*. The resultant platelet-containing pellet was resuspended in 10 mM HEPES (pH 7.4), 134 mM NaCl, 12 mM NaHCO_3_, 2.9 mM KCl, 0.34 mM Na_2_HPO_4_, and 1 mM MgCl_2_ (Tyrode’s buffer) containing 0.5 μM prostaglandin (PGI_2_). Platelets were then washed in PGI_2_-free Tyrode’s buffer, containing 5 mM dextrose and 0.3% BSA, and counted in a haemocytometer.

### Flow cytometry

Washed platelets (1.5 × 10^5^ platelets/μl) were maintained in Tyrode’s buffer (unactivated) or treated with 30 μM ADP (activated). Both unactivated and activated platelets were incubated for 6 min at room temperature with either control liposomes (Lipo-C, 50 μg/ml) or sulfatide-containing liposomes (Lipo-S, 50 μg/ml) in the absence and presence of 50 μM of one of the following peptides: Dab2 SBP, Dab2 SBP R42A, or Dab2 SBP R42K. Platelets were fixed with 1% formalin in phosphate buffered saline and incubated with phycoerythrin-labelled anti-CD62P (P-selectin; Biolegend) antibody following manufacturer’s instructions. Antibody-bound platelets were quantified using a FacsAria flow cytometer.

## RESULTS

### Refining the sulfatide-binding site in Dab2 SBP

Previous NMR studies showed that the last 20 residues of Dab2 SBP, with a C-terminal polybasic region 49-KYKAKL-54 (**Fig. 1A**), plays a major role in sulfatide interactions (17). Molecular docking was performed to refine potential residue interactions between a single sulfatide molecule and the model of the DPC-bound structural conformation of Dab2 SBP (PDB ID: 2LSW) containing vector residues (19-GPLGS-23) added to the N-terminus to match the experimental peptide used in this work (17). After energy minimization, the structure of Dab2 SBP was validated and showed favorable energetics and side chain positioning using Ramachandran plot, ProSA and QMEAN analyses (**Fig. S1**). All nine docking poses hit a similar Dab2 SBP scaffold and indicated that P32 (backbone), K30 (backbone), and R42 residues form hydrogen bonds with the OS atoms of the sulfate group in the sulfatide, whereas the acyl chains faced the α-helix 2 of the peptide (**Fig. 1B-C**). Specifically, the side chain of R42 interacted with the sulfatide head group forming hydrogen bonds and electrostatic interactions that were approximately 3Å apart (**Fig. 1C**). Both E33 and Y38 interacted with the galactose moiety (**Fig. 1C**). The conserved 49-KYKAKL-54 region in Dab2 SBP (**Fig. 1A-C**) was initially suggested to provide an electrostatic environment to attract and accommodate the negatively charged sulfatide in its Dab2 SBP binding pocket. However, molecular docking results showed a strong R42 interaction with the sulfate head group, which led to consistent positioning of the acyl tails of sulfatide towards the α-helix 2. Initial docking results and fingerprinting showed that the CH2 moieties of the K49 and K51 residues interacted with the acyl tails. As K49 and K51 are critical for sulfatide binding (13), the docking results suggest that the four CH2 moieties in K49 and K51 contribute to a hydrophobic microenvironment (**Fig. 1D**) in the α-helix 2 of Dab2 SBP, which aid in accommodating acyl tails and leading to more energetically favorable positioning of the sulfate head group towards R42 (**Table S1**). Hydrophobic interactions in α-helix 2 were also provided by Dab2 SBP residues Y50, L54, and I55 (**Fig. 1C and Table S1**). Altogether, molecular docking data indicate that α-helix 1 of Dab2 SBP is involved in contacts with the sulfatide head group, whereas α-helix 2 favors hydrophobic contacts with the acyl chains of the lipid.

**Figure 1.**
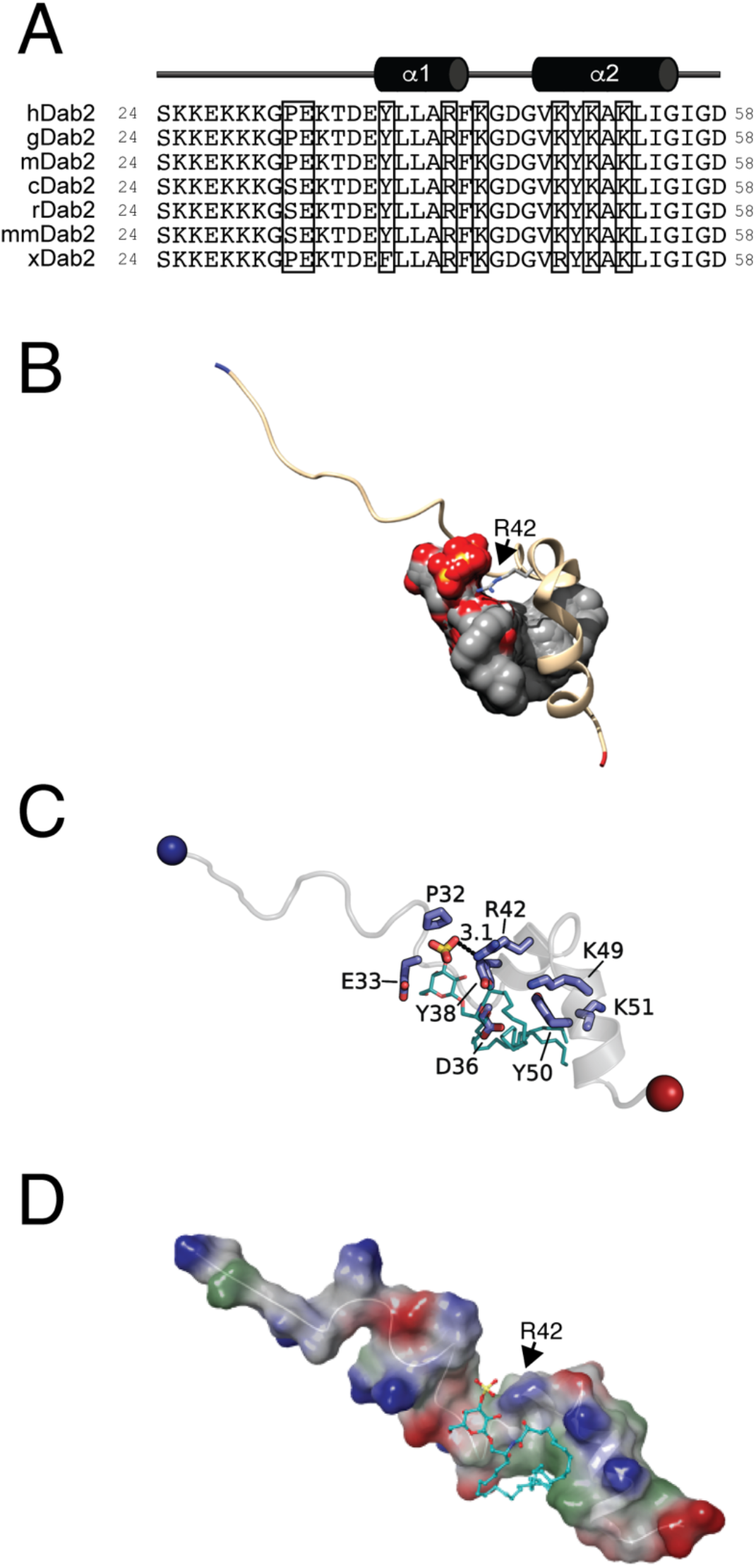
Sulfatide docking onto Dab2 SBP. (**A**) Sequences of the closest homologues of Dab2 proteins corresponding to the SBP region. hDab2, *Homo sapiens* Dab2; gDab2, *Gorilla gorilla* Dab2; mDab2, *Mandrillus leucopheous* Dab2; cDab2, *Canis lupus familiaris* Dab2; rDab2, *Rattus norvegicus* Dab2; mmDab2, *Mus musculus* Dab2; xDab2, *Xenopus laevis* Dab2. Residues implicated in sulfatide binding, as determined from this work, are boxed. (**B**) Overlaid poses of sulfatide docked to Dab2 SBP. Dab2 SBP is rendered as a cartoon and is colored tan with the N-terminus colored blue and the C-terminus is colored red. R42 is shown as a stick that is colored gray and by atom type. The nine poses produced by AutoDock Vina are shown as a gray surface and by atom type. The side chain of R42 is colored in blue stick. Sulfatides (cyan) are shown as sticks and colored by element. (**C**) Key sulfatide-binding residues of Dab2. Dab2 SBP is rendered as a cartoon colored transparent gray with the N- and C-terminus shown as blue and red spheres, respectively. Key residues are shown as blue sticks and labelled. (**D**) Surface representation of sulfatide-bound Dab2 SBP showing the hydrophobic (green), positively charged (blue), and negatively charged (red) surface regions. Sulfatide backbone is represented in stick colored with carbon as cyan, sulfate as yellow, and oxygen as red. Surface potential was calculated using Schrodinger Maestro.

### Conformational flexibility of Dab2 SBP upon sulfatide binding

In agreement with our previous work (17), addition of DPC-embedded sulfatide to DPC-containing Dab2 SBP has little or no effect on ^1^H and ^15^N chemical shifts of residues S24-E37 but perturbs resonances of most of the residues spanning residues Y38 to D58 (**Fig. S2A**). The heights of HSQC peaks for residues E33-I56 are considerably lower than those for residues S24 -G31 for both DPC-containing Dab2 SBP with and without sulfatide-embedded micelles (**Fig. S2B**). Residues S24-G31 do not likely contact DPC micelles and are highly mobile and solvent-exposed as suggested from paramagnetic relaxation experiments (17). Residues Y38-I55, on the other hand, contribute to the secondary structure in Dab2 SBP and strongly interact with DPC micelles (17). Consequently, as observed in **Fig. S2B**, residues E33-I56 may be poorly observable with solution NMR when Dab2 SBP is bound to either sulfatide-free or -embedded DPC micelles.

We also used NMR to characterize the backbone dynamics of Dab2 SBP in its free-and sulfatide-bound states (**Fig. 2**). Picosecond to nanosecond backbone dynamics, measured by ^1^H–^15^N, heteronuclear Overhauser effects (NOE) of DPC-associated Dab2 SBP, showed very flexible N- and C-termini and a rigid structure spanning residues E37-I56; such flexibility was not altered by sulfatides (**Fig. 2A**). Addition of sulfatides decreased *R*_1_ and increased *R*_2_ of residues T35-I56. This suggests that the correlation time of the motions causing relaxation is greater than the molecular correlation time at the *R*_1_ maximum (~ 5×10^−8^ s) of a *R*_1_ versus the molecular correlation time plot (30). The relaxation data also indicate that sulfatide has little or no effect on the motions of residues S24-G31 but shifts the spectral density of residues T35-I56 to a lower frequency. This could be due to sulfatide binding to residues T35-I56, consistent with the data from both molecular docking (**Fig. 1B-C**) and HSQC titrations of the peptide with the lipid (**Fig. S2**), or a sulfatide-induced increase in DPC micelle size or both.

**Figure 2.**
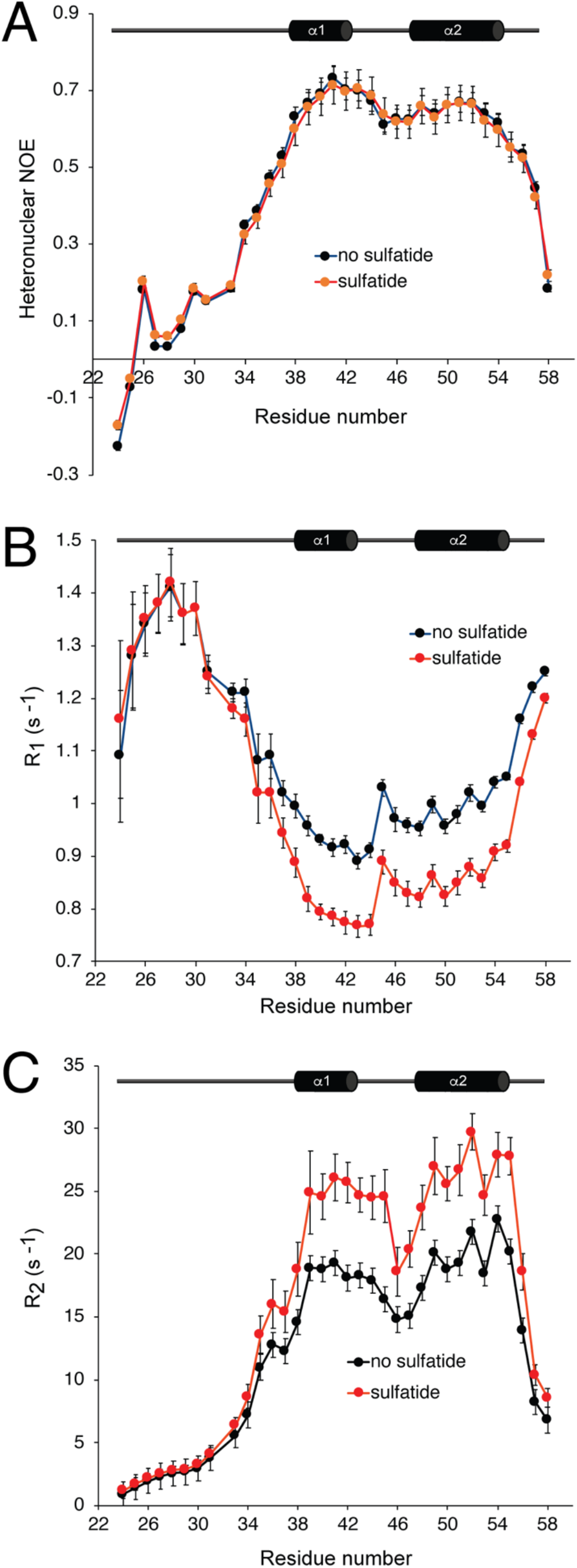
Dab2 SBP dynamics characterized by NMR relaxation measurements. ^1^H–^15^N NOE ratio (**A**), longitudinal relaxation rates, *R*_*1*_ (**B**), and transversal relaxation rates, *R*_*2*_ (**C**) of DPC-embedded Dab2 SBP in the absence (black) or presence (red) of 8-fold DPC-embedded sulfatides. The secondary structure of Dab2 SBP is depicted at the top of each panel.

### *In silico* and lipid-binding assays of Dab2 SBP mutants confirm critical sulfatide-binding residues

From sulfatide docking studies on wild-type Dab2 SBP, sulfatide docked 8 out of 9 poses clustered near R42 with the sulfate groups being within 5.6 Å between the furthest atoms of the sulfate group in each pose and 1 out of 9 poses residing within 7.4 Å of the cluster group. In addition, HSQC titrations showed that sulfatide markedly perturbed R42 (**Fig. S2A**), consistent with previous observations (17). Further molecular docking experiments were performed to confirm the influence of R42 for sulfatide binding. Replacement of R42 to alanine reduced close contacts of Dab2 SBP for sulfatide, as concluded from the analysis of nine independent sulfatide poses on the mutated peptide (**Fig. 3A**, **Fig. S3**, and **Fig. S8**). Dab2 SBP R42A increased the distance for sulfate head group-A42 interactions to greater than 5.0 Å compared to the wild-type peptide (3.1 Å). To adjust to the loss of R42 in Dab2 SBP R42A, the sulfate head group sought to hydrogen bond with E33 and T35. Interaction fingerprint analysis of Dab2 SBP R42A showed fewer overall interactions (**Fig. 3A**), specifically a loss of aromatic and polar interactions with sulfatide compared to the wild-type peptide (**Table S1**). Docking results also revealed that a mutation of R42 to lysine (R42K) reduced the number of Dab2 SBP contacts with sulfatide (**Fig. 3B**) and exhibited less specific docking clusters on the predicted sulfatide-binding site in Dab2 SBP (**Fig. S8**), but more closely resembled wild-type Dab2 SBP interactions than R42A. Further analysis indicated that R42K had less aromatic, charged, hydrophobic, and polar interactions while having more hydrogen bond interactions compared to Dab2 SBP (**Table S1**). Although Dab2 SBP R42K retains the charged interaction that R42 displayed in Dab2 SBP, the interaction distance between the side chain of K42 and the sulfate head group increased to 3.9 Å. This result indicates that not only the charged but also the stereochemistry of an arginine residue is required for proper sulfatide docking in Dab2 SBP. Similarly, replacement of Y38 to alanine led Dab2 SBP to have less polar, hydrophobic, and charged interactions with sulfatide while having more backbone and hydrogen bond interactions (**Table S1**). Dab2 SBP Y38A increased the interaction distance to 5.0 Å with the sulfate head group (**Fig. 3C**), confirming that Y38 plays an important role in sulfatide docking, while not necessarily interacting with the sulfate group itself.

**Figure 3.**
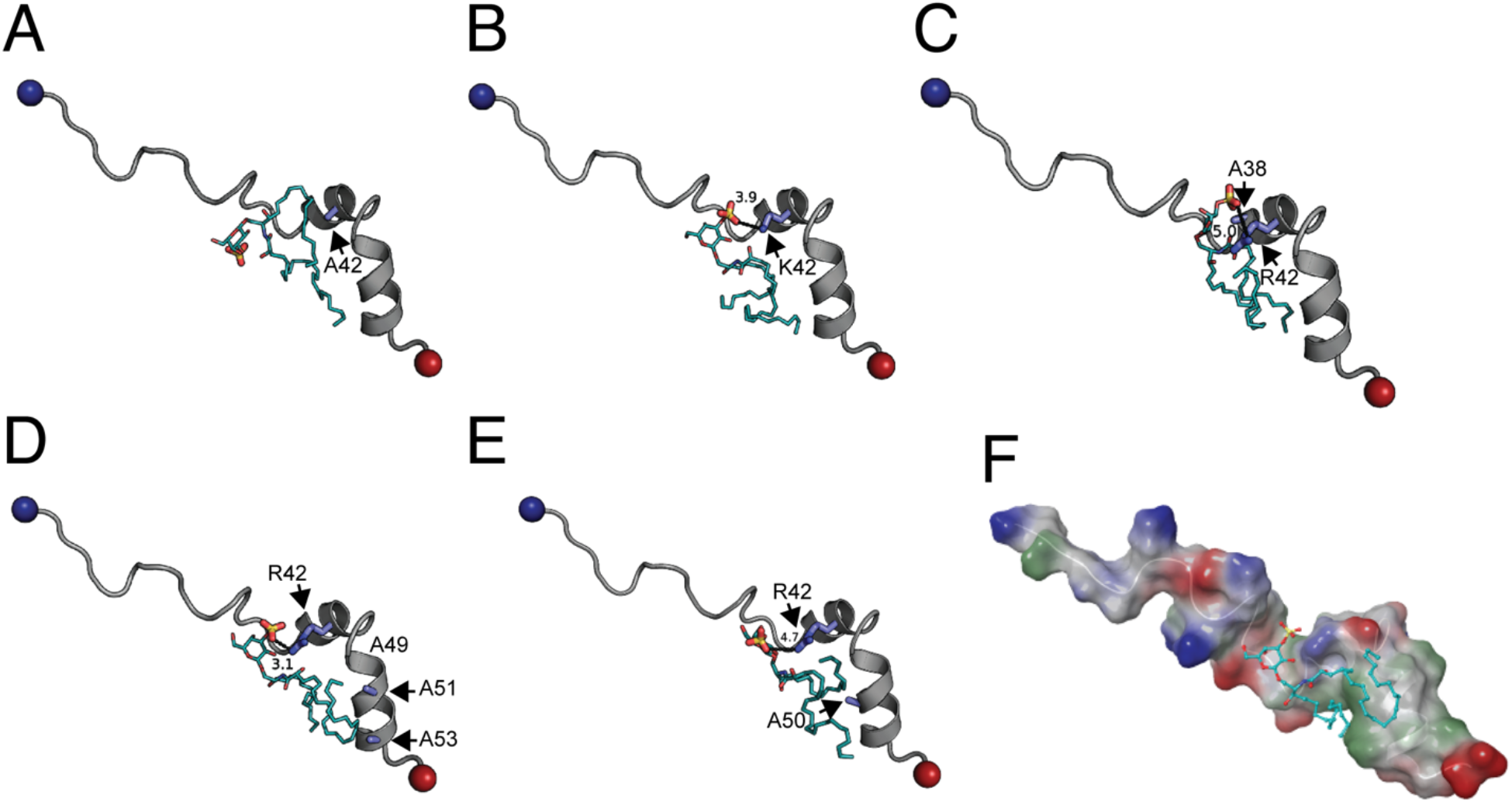
Identification of Dab2 SBP critical residues for sulfatide binding using molecular docking. (**A-E**) Lowest energy pose of sulfatide docked to Dab2 SBP R42A (**A**), Dab2 SBP R42K (**B**), Dab2 SBP Y38A (**C**), Dab2 SBP K49A/K51A/K53A (**D**), and Dab2 SBP Y50A (**E**). In all cases, key residues are colored in blue. Sulfatides (cyan) are shown as sticks and colored by element. (**F**) Surface representation of sulfatide-bound Dab2 SBP K49A/K51A/K53A showing the hydrophobic (green), positively charged (blue), and negatively charged (red) regions. Sulfatide backbone is represented in stick colored with carbon as cyan, sulfate as yellow and oxygen as red. Surface potential was calculated using Schrodinger Maestro.

Molecular docking of sulfatide in Dab2 SBP predicted lipid acyl chain interactions with residues located in α-helix 2 (**Fig. 3D-E**). Combination mutations at K49, K51, and K53 stabilized the R42 and sulfate head group interaction but influenced acyl tail positioning (**Fig. 3D** and **Fig. S6**). This triple mutation in Dab2 SBP showed the same interaction distance as the wild-type peptide (3.1 Å) with tight sulfate head clustering, as all 9 poses were clustered within 5.5 Å of each other and interacted with R42 compared to 8 out of 9 poses in the wild-type form (**Fig. S8**). However, Dab2 SBP K49A/K51A/K53A had less polar, aromatic, and charged interactions while having more backbone and hydrogen bond interactions (**Table S1**). The surface rendering of Dab2 SBP K49A/K51A/K53A indicated an exposed hydrophobic area that is less delimited than the wild type Dab2 SBP (**Fig. 3F**), potentially causing a loss of affinity for sulfatide. Thus, these results support the observed environment of Dab2 SBP, generated by residues R42, Y38, K49, K51, and K53, for sulfatide binding. Replacement of Y50 by alanine in the peptide exhibited less total charged and hydrophobic interactions with sulfatide, increasing the sulfate head group distance to R42 to 4.7 Å with a concomitant increment in polar, backbone, and hydrogen bond interactions (**Fig. 3E**, **Fig. S7,** and **Table S1**).

To experimentally assess whether the mutations in Dab2 SBP alter sulfatide binding, recombinant Dab2 SBP, and the mutants identified by docking studies, were fused to GST and employed to screen for sulfatide binding using the lipid-protein overlay assay. Although this assay does not mirror the physiological organization of sulfatides in biological membranes, it is still useful to obtain a first screening for the binding of the Dab2 SBP mutants to the sphingolipid. As expected, Dab2 SBP bound sulfatides and closely reached saturation at 800 pmoles of lipid (**Fig. 4A-B**). Alanine mutations at residues upstream of α-helix 1, such as P32 and E33 (**Fig. 1A**), reduced the peptide’s sulfatide binding, whereas mutation at R42 abolished it (**Fig. 4**). The poor binding capacity to sulfatides displayed by Dab2 SBP R42A is not due to a disruption of α-helix 1 as it remains as folded as the wild-type peptide, as indicated by their circular dichroism spectra (**Fig. S9**). Consistent with the molecular docking studies, R42 to lysine reduced sulfatide binding whereas alanine mutations on residues outside of the predicted binding pocket, such as E37 and E46, did not alter lipid binding (**Fig. 4**). Unexpectedly, alanine mutation of Dab2 SBP at K44, which did not alter the secondary structure of the peptide (**Fig. S9**), markedly affected sulfatide binding (**Fig. 4**). Mutagenesis at the second helix, where sulfatide acyl chains may interact, showed that Y50 is not critical for sulfatide binding, but a triple alanine mutation at residues K49, K51, and K53 markedly reduced it (**Fig. 4**) without affecting the secondary structure of the peptide (**Fig. S9**). The role of these basic residues in sulfatide-binding is in agreement with mutagenesis and liposome-binding studies we obtained previously using Dab2 N-PTB (13).

**Figure 4.**
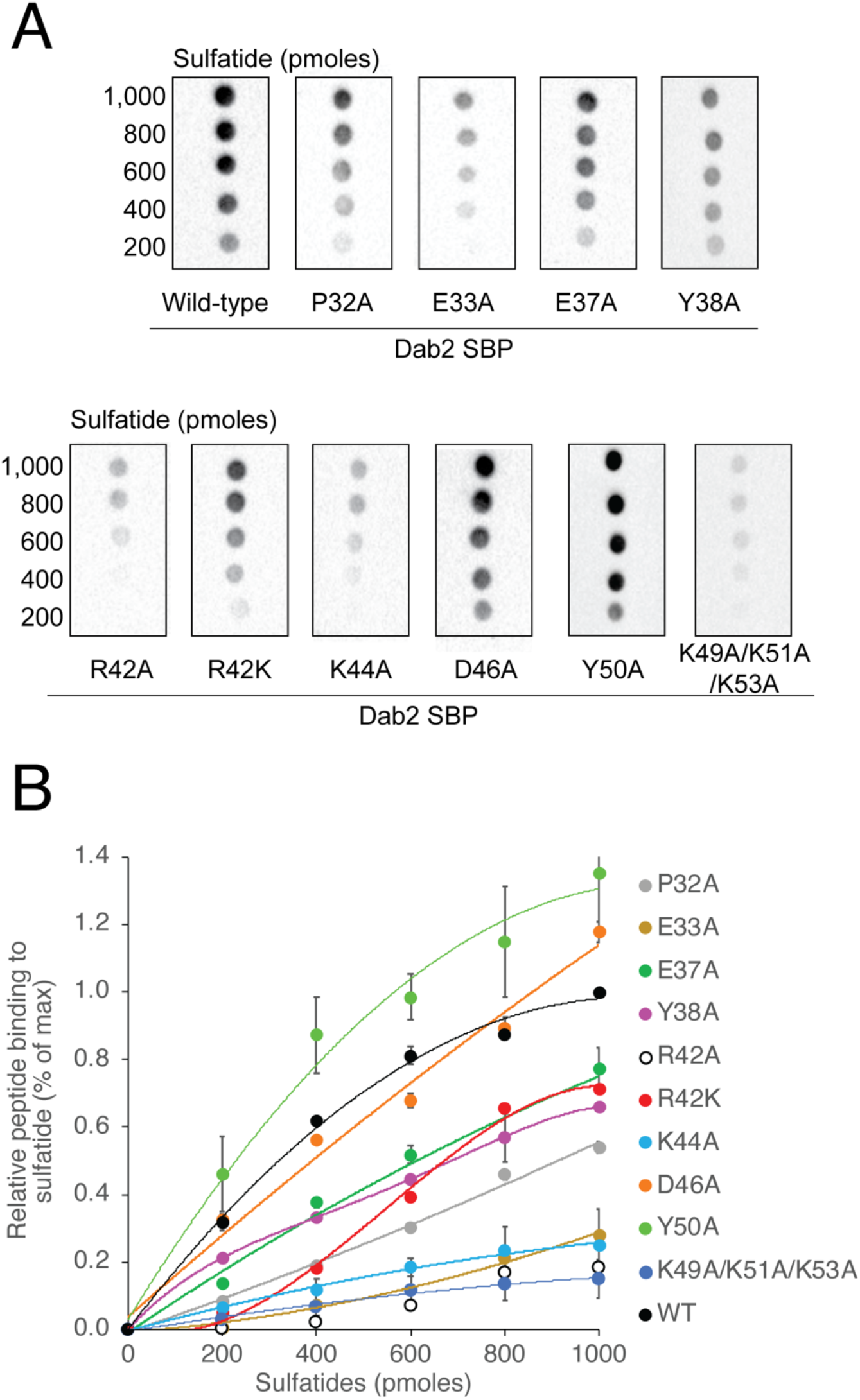
Characterization of sulfatide binding to Dab2 SBP *in vitro*. (**A**) Lipid-protein overlay assay displaying the binding of GST fusion Dab2 SBP and the indicated mutants to sulfatides immobilized on nitrocellulose. (**B**) Quantification of the binding of the GST fusion peptides to sulfatides.

Next, taken the molecular docking and LPOA results together, we focused on the novel role of the Dab2 SBP residues Y38 and R42 for binding to sulfatides using sulfatide-enriched liposomes of 100 nm in diameter and followed their interactions using SPR. In agreement with previous observations (17), SPR data showed that Dab2 SBP bound sulfatide liposomes with a *K*_D_ of ~35 μM and the association displayed saturation of binding (**Fig. 5A** and **D**). Alanine mutation at Y38 exhibited a decrease of sulfatide liposome binding with a *K*_D_ of ~70 μM (**Fig. 5B** and **D**). Binding of Dab2 SBP R42A to sulfatide liposomes was weaker, with an estimated *K*_D_ higher than 100 μM (**Fig. 5C-D**). Thus, in addition of the role of the α-helix-2 in sulfatide binding, our results show that R42 and Y38, the latter in a lesser extent, are relevant for the association of Dab2 SBP with sulfatide-embedded lipid bilayers.

**Figure 5.**
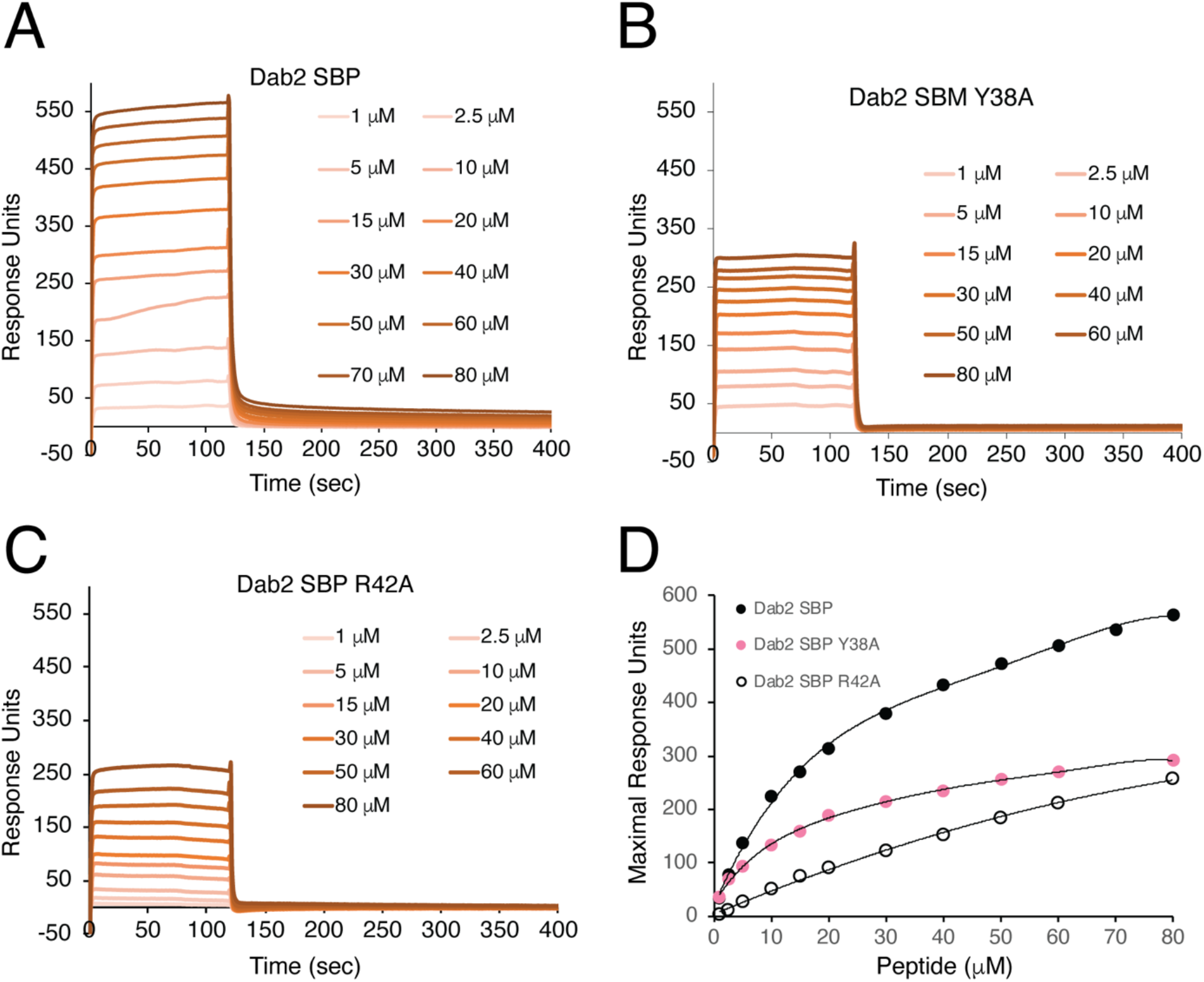
Critical Dab2 SBP residues for binding of mimics of sulfatide-containing lipid bilayers. (**A-C**) SPR traces depicting the binding of Dab2 SBP (**A**), Dab2 SBP Y38A (**B**), and Dab2 SBP R42A (**C**) from 1 to 80 μM, to sulfatide liposomes. (**D**) Plot representing the binding of Dab2 SBP (black circles), Dab2 Y38A (pink circles), and Dab2 SBP R42A (empty circles), from 1 to 80 μM, to sulfatide-containing liposomes.

### Dab2 SBP depends on R42 to reduce sulfatide-mediated P-selectin expression

P-selectin is a cell surface pro-aggregatory protein found in platelets and endothelial cells. This protein is stored in α-granules and is quickly released to the platelet surface in response to a wide range of thrombogenic stimuli (31). P-selectin promotes platelet aggregation in a sulfatide-dependent manner (32). To establish whether Dab2 SBP can modulate P-selectin pro-aggregatory activity, we evaluated the cell surface expression of P-selectin. Washed platelets were previously activated with ADP, which represents a critical step for sulfatide-mediated platelet activation (32). Consistent with previous observations (15), incubation of activated platelets with sulfatide-containing liposomes led to an increase in the median fluorescence of the anti-P-selectin antibody bound to the platelet surface (**Fig. 6A**), indicating that sulfatides promote P-selectin accumulation at the platelet surface. Pre-incubation of sulfatide liposomes with Dab2 SBP significantly reduced the presence of P-selectin at the platelet surface (**Fig. 6A**), suggesting that the peptide competes with P-selectin for sulfatide binding. Replacement of R42 with lysine or alanine significantly impaired the Dab2 SBP inhibitory activity (**Fig. 6A**). Representative histograms showing the distribution of the baseline population of platelets is indicated in **Fig. 6B**. As expected, ADP induces a minor right shift, which is indicative of platelet activation (**Fig. 6B**, inset). In contrast to sulfatide-free liposomes, the majority of the platelet population become further activated by the presence of sulfatide liposomes (**Fig. 6B**, inset). In contrast to the presence of either R42A or R42K, the addition of Dab2 SBP led to a shift to the left of the median fluorescence (**Fig. 6B**). Together, our results indicate that Dab2 SBP has the potential to inhibit P-selectin function, with R42 playing a critical role.

**Figure 6.**
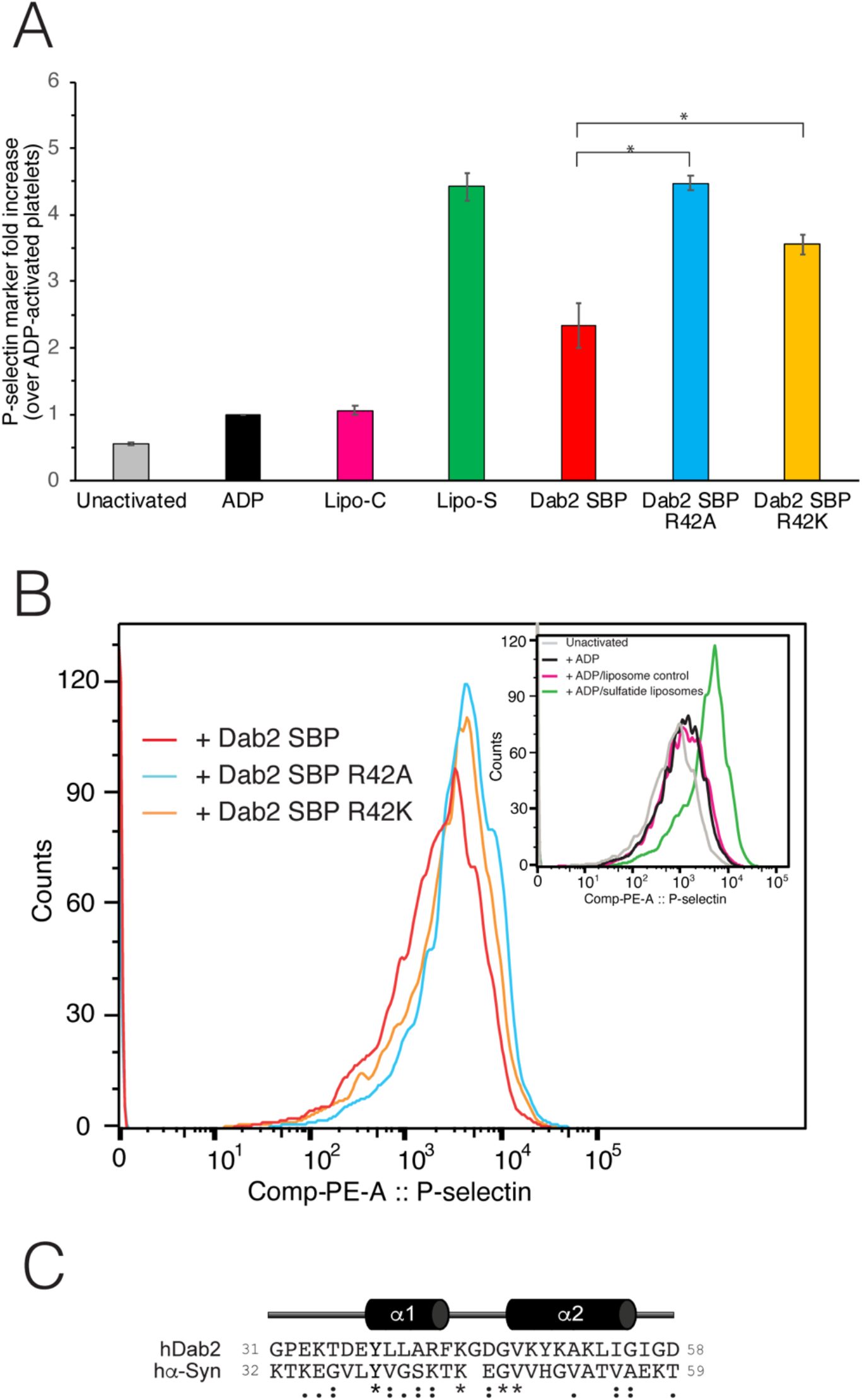
Inhibition of sulfatide-induced platelet P-selectin surface expression in platelets by Dab2 SBP. (**A**) Platelets were stimulated with ADP and further incubated with sulfatide-free or -containing liposomes in the absence or presence of the indicated peptides. Samples then were fixed, incubated with PE-labeled CD62P (anti-P-selectin) antibody, and analyzed by flow cytometry. The graph represents the median fluorescence intensity for each treatment (mean ± standard deviation) of three independent experiments. Data is represented as a fold increase in fluorescence over ADP-treated platelets. Statistical analysis was carried out using a *t*-test. (**B**) Color-coded representative immunofluorescence histogram displaying the presence of platelet surface P-selectin for the treatments indicated in **A**. The black plot in the inset indicates the presence of P-selectin in the surface of unactivated platelets. (**C**) Comparison of the α-synuclein sphingolipid-binding domain with the sulfatide-binding site of Dab2. Asterisks represent identical residues, whereas residues that share common properties are shown as colons. Per ClustalW criterium, semiconservative substitutions are indicated with dots.

## DISCUSSION

In this study, we structurally and functionally characterized the sulfatide binding region of Dab2, represented by SBP (human Dab2 residues 24-58). Our molecular docking studies suggest that residues upstream and on the first α-helix of Dab2 SBP interact with the sulfatide head group, whereas the second α-helix provided recognition of the sphingolipid acyl chains. To establish the structural basis of Dab2 interaction with sulfatides, we designed a series of Dab2 SBP mutants, which were tested for *in vitro* sulfatide binding. In comparison to our previous report using Dab2 N-PTB (13), the N-terminal 25-KKEKKK-30 region in Dab2 did not exhibit a role in sulfatide binding as observed from our molecular docking and NMR relaxation results.

Our current results suggest that the Dab2 SBP basic residue, R42, is critical for association to the negatively charged sulfatide. *In silico* results closely matched our experimental results, highlighting the ability of docking to predict key residues for binding sulfatide in Dab2 SBP. Replacement of R42 with lysine reduced sulfatide binding *in vitro* and significantly affected the function of the peptide for targeting P-selectin surface expression in platelets. This result implies that not only the positive charge, but also the stereochemistry of the side chain, is required for sulfatide interactions. In addition, other charged and aromatic Dab2 SBP residues (*i.e*., E33, Y38, K44) were relevant for lipid interactions. Previous results suggested that the last 20 residues of Dab2 SBP, containing residues Y38-L40 and R42 as well as the C-terminal 49-KYKAKL-54 motif are involved in sulfatide interactions (17). Mutagenesis analysis of the C-terminal 49-KYKAKL-54 motif (**Fig. 4**) confirmed its role in sulfatide binding. By docking a sulfatide to Dab2 SBP, we found that the C-terminal polybasic motif, located in α-helix 2, is likely required for acyl chain hydrophobic interactions rather than binding to the sphingolipid head group. Lysine residues are considered to exhibit a dual role in lipid contacts, that is, by promoting electrostatic interactions with negatively charged lipids and/or by employing their flexible hydrocarbon spacers for hydrophobic interactions with membrane lipids (33). Indeed, we observe a hydrophobic patch in the wild type Dab2 SBP structure (**Fig. 1D**) that is shaped by K49, K51, and K53. The suggested roles of the α-helices in sulfatide docking are in agreement with the doxyl stearic acid-based paramagnetic quenching NMR experiments using sulfatide-embedded DPC micelles (17), which demonstrate that the first Dab2 SBP α-helix is less quenched than the second one. Thus, this suggests that the first α-helix is closer to the micellar surface, whereas the second α-helix is oriented towards the micelle core.

Very few protein-sulfatide complexes have been structurally characterized. The interaction of the cluster of differentiation 1a (CD1a) with sulfatide involves hydrogen bonds to its 3’ sulfate group with R76 and E154 and the galactose moiety with R76 and S77, whereas the sulfatide fatty acid chains display van der Waals interactions mostly with hydrophobic residues (34). Interestingly, a basic residue (H38) plays a role in accommodating one of the alkyl chains in the binding pocket of CD1a.

Sphingolipid-binding domains, such as those described in viral, bacterial, and mammalian proteins (35–37), exhibit a consensus helix-turn-helix fold with an aromatic residue, predominantly, being solvent-exposed, and either a G or P residue contributing to the turn between the α-helices and several positively charged residues (38). Intriguingly, a consensus sphingolipid-binding motif has been identified in α-synuclein (39), which shares several features with Dab2 SBP (**Fig. 6C**). Molecular dynamic simulations show that α-synuclein simultaneously interacts with two molecules of the glycosphingolipid monosialodihexosylganglioside (GM3), with Y39 representing the most critical sphingolipid-binding residue (39). Of note, it is currently unknown whether Dab2 can bind two sulfatide molecules simultaneously. The α-synuclein protein also binds sulfatides, but GM3 represents its preferred ligand. Indeed, we showed that alanine mutation in the equivalent conserved tyrosine residue in Dab2, Y38, reduces the affinity for sulfatide binding by ~50%. In the α-synuclein membrane model interaction, it is proposed that Y39 is located at the interface between the polar and nonpolar regions of the sphingolipid, mediating the protein insertion into the membrane (39). Supportive of this observation, Dab2 Y38 undergoes 30-50% reduction in its resonance intensity in micellar paramagnetic NMR experiments (17). Accordingly, we observed that Dab2 SBP Y38 exhibited minor dynamic changes when the peptide was in contact with sulfatide-embedded DPC micelles. A basic residue (K34) located upstream of Y39, also displays a major role for α-synuclein interactions with GM3 (39). However, Dab2 SBP K34 did not exhibit an important role in sulfatide binding as concluded from molecular docking and NMR assays. In contrast to GM3, sulfatides are negatively charged. Thus, it is possible that α-synuclein K43 may play a similar role to that observed for Dab2 R42 in sulfatide interactions. Considering that R42K mutation in Dab2 SBP reduced the activity of the peptide but did not abolish it, the presence of K43, instead of R43, might also reduce the preference of α-synuclein for sulfatides.

Sulfatides are found in discrete patches on activated spread platelet membranes, reminiscent of lipid rafts. Platelet surface expression of the sulfatide-binding pro-aggregatory protein P-selectin increases the stability of platelet aggregation (32). Accordingly, Dab2 modulates the extent of platelet aggregation by targeting P-selectin function. The association of Dab2 to sulfatides impairs P-selectin-sulfatide interactions, thereby blocking homotypical and heterotypical cell-cell interactions (15). Upon platelet activation, Dab2 is transiently secreted from α-granules to the cell surface (13). Upon sulfatide binding, Dab2 likely undergoes a conformational change (40), a state in which the protein is more tolerant to thrombin proteolysis (13). It has been suggested that, once secreted, Dab2 modulates the extent of platelet aggregation by two independent mechanisms (1). First, Dab2 downregulates the integrin receptor activity by interfering with its interaction with fibrinogen. Second, extracellular Dab2 blocks platelet-platelet P-selectin-mediated sulfatide-dependent interactions. Other sulfatide-binding proteins have also been reported to be exported out of the cell for extracellular interactions with sulfatides. For example, the mesencephalic astrocyte-derived neurotrophic factor (MANF) is an endoplasmic reticulum protein that is secreted under stress-related stimuli (41). Binding of secreted MANF to sulfatides, through its α-helical saposin-like domain, provides MANF cellular uptake and alleviates stress responses driven by the endoplasmic reticulum (42).

In conclusion, our results indicate how sulfatides interact with Dab2 SBP and identify the critical binding residues. Our data is in line with what was earlier defined as the minimal α-helical-turn-α-helical unit for sphingolipid binding. Furthermore, in the case of Dab2, R42 mediating the recognition of negatively charged sulfatide headgroup, and the α-helical lysine residues providing an adequate hydrophobic environment through their flexible hydrocarbon spacers.

## Abbreviations

CD: circular dichroism
Dab2: Disabled-2
DPC: dodecylphosphocholine
GST: glutathione s-transferase
N-PTB: N-terminus containing the phosphotyrosine-binding domain
SBP: sulfatide-binding peptide
SPR: surface plasmon resonance.

## Acknowledgements

We thank Dr. Janet Webster for critical reading of the manuscript. This project was supported by the Virginia Academy of Science, the Institute for Critical Technology and Applied Science (ICTAS) at Virginia Tech, and the 4-VA Collaborative Research Program (to D.G.S.C.) and the National Science Foundation (MCB-1517298) (to C.V.F). W.S. was supported by an ICTAS pre-doctoral fellowship.

